# SoftWipe – a tool and benchmark to assess scientific software quality

**DOI:** 10.1101/2020.10.07.330621

**Authors:** Adrian Zapletal, Dimitri Höhler, Carsten Sinz, Alexandros Stamatakis

**Author notes:** equal contribution (Adrian Zapletal, Dimitri Höhler).

## Abstract

Scientific software from all areas of scientific research is pivotal to obtaining novel insights. Yet the quality of scientific software is rarely assessed, even though it might lead to incorrect scientific results in the worst case. Therefore, we have developed an open source tool and benchmark called `SoftWipe`, that provides a relative software quality ranking of 51 computational tools from diverse research areas. `SoftWipe` can be used in the review process of software papers and to inform the scientific software selection process.

---

Scientific software (henceforth called software) has become pivotal for discoveries in almost all research fields, including emerging areas such as the digital humanities. The importance of software has become evident to the broad public by the plethora of computational tools (henceforth called tools) used to analyze SARS-CoV-2 data, for instance. However, due to the conditions under which software is typically being developed, such as, lack of sustainable funding for maintaining widely used tools, lack of time, and a lack of formal training in computer programming and software engineering, software quality exhibits a large variance. More importantly, software quality is typically not taken into account in the software development, software paper review, or in the tool selection process, for instance, when two computational tools provide analogous functionality. This is particularly worrisome as political decisions are currently partially based on such tools for handling the SARS-CoV-2 pandemic.

The term ’software quality’ is admittedly fuzzy, but offers a handle to perform largely automated analyses of scientific open source tools based on a set of objective criteria and measures. Such an automated analysis of numerous tools would be hard if one wanted to apply proper software verification techniques. When a tool exhibits ’bad’ software quality, this does absolutely not imply that it is erroneous. However, it does imply that the probability for it to generate faults that might, in the worst case, lead to paper retractions or incorrect results obtained with the faulty tool is higher (based on results from the area of empirical software engineering (1, 2)). See the introduction of (3) for a list of some high profile retractions and corrections because of software errors. For the above reasons, we strongly advocate that software quality should be assessed in the development, review, and tool selection process.

To this end, and based on our previous work (4) where we manually assessed the software quality of widely used tools for evolutionary biology, we introduce `SoftWipe`, a largely automated open source tool and benchmark for assessing the software quality of open source tools written in `C` or `C++`. We envision that `SoftWipe` can be used by authors/developers (e.g., requiring them to provide their `SoftWipe` score with the submission), reviewers and researchers to integrate software quality into the review process and to raise awareness about software quality and engineering issues which are crucial for the success of computational science in general.

The `SoftWipe` score is a relative score over a number of software quality indicators (see further below and the supplement for a description of these indicators). The best-ranking tool per indicator receives a score of 10 while the worst ranking tool per indicator receives a score of 0. Subsequently, we compute the average unweighted score per tool over all per indicator scores to obtain a global software quality ranking.

This global relative score might change over time as more tools are being added to the benchmark and will therefore be difficult to reference. We have therefore also devised an absolute fixed global score (see supplement for mathematical details) that does not change when further tools are added to the benchmark and that can therefore easily be referenced.

We use the following software quality indicators (normalized by average values per 1000 lines of code) to rate the tools: number of compiler, sanitizer, and static code analyzer warnings as generated by a variety of tools, number of assertions used, cyclomatic code complexity, indicators of bad programming style, and degree of code duplication. A detailed description of the quality indicators including a rationale for the inclusion of each one is provided in the supplement.

The current benchmark comprises 51 tools from a wide range of scientific fields such as, evolutionary biology, astrophysics, epidemiological simulation, hypergraph analysis, deep learning applied to DNA data, antigen typing, protein data structure mining, SAT (satisfiability) solvers, etc. The benchmark results are provided in Table 1 Note again that absolute fixed `SoftWipe` scores do not change when new tools are included in the benchmark, while relative scores do change (see supplement for details). A comprehensive table including all individual software quality indicators as well as descriptions of the respective tools is available online at https://github.com/adrianzap/SoftWipe/wiki/Code-Quality-Benchmark.

**Table 1.**
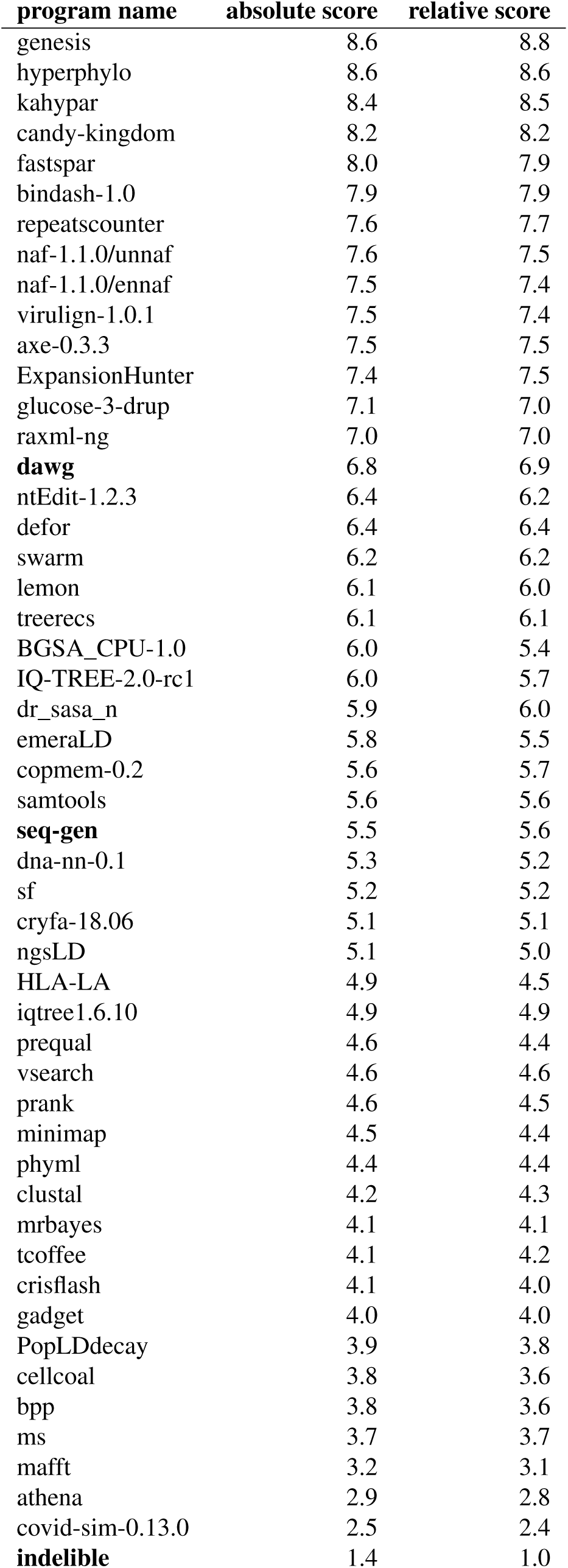
Absolute fixed and relative `SoftWipe` scores of 51 computational tools from a wide range of scientific fields such as computer science, evolutionary biology, astrophysics, and bioinformatics. The three DNA sequence simulation tools dawg, seq-gen, and indelible that have similar functionality are highlighted in bold font.

One important observation is that the four top scoring tools have all been developed by researchers with formal training in pure computer science whereas the 10 worst scoring tools have all been developed by self-taught programmers without formal training in computer science.

For an example of how the benchmark can be used, let us consider the three evolutionary molecular sequence data simulators `dawg, seq-gen`, and `indelible` (highlighted in bold font in Table 1). These tools implement highly similar functionalities and models. Our benchmark clearly suggests the use of `dawg`, as it obtains a higher score than the alternative sequence simulators.

It is also noteworthy that the developers of the top three tools had access to a pre-release version of `SoftWipe` as they are members or collaborators of our lab and improved their tools to obtain higher `SoftWipe` scores. This highlights the utility of `SoftWipe` within a lab such as ours that predominantly develops computational tools, as it fosters a healthy competition among lab members to develop the best-scoring tool. In fact, the regular deployment and integration of `SoftWipe` in the development process has also helped to avoid a bug in at least one occasion.

Given the current SARS-CoV-2 pandemic it is also worth noting that the `CovidSim` tool obtains the second worst score among all 51 tools tested.

We are well aware of the fact that `SoftWipe` is likely to generate controversial debates, as software quality does neither induce lack of code correctness, nor an erroneous underlying scientific model. Nonetheless, in the current absence of any standard routines for assessing scientific software quality and based upon the results of empirical software engineering research that establishes a correlation between software quality and failure, `SoftWipe` represents a step forward, as it is easy to use and does not require a substantial investment of time. This does not only apply to the software submission and review process, but also to software development. `SoftWipe` can also help to objectify controversial discussions about software quality, as was the case with the CovidSim tool (see e.g., https://www.imperial.ac.uk/news/197875/codecheckconfirms-reproducibility-covid-19-model-results). We are therefore convinced that a routine deployment of `SoftWipe` or analogous tools at various stages of the software development and publication process can substantially contribute to improving software quality across all scientific fields, raise general awareness about the importance of software quality in the respective research communities, and reduce the number of program faults.

## Methods

The open source code of `SoftWipe` including the up-to-date benchmark and a user manual is available at: https://github.com/adrianzap/SoftWipe/wiki.

## Acknowledgements

Part of this work was funded by the Klaus Tschira foundation. We wish to thank Bernd Doser for initial discussions on `SoftWipe` and Frédéric Mahé for useful comments on this manuscript.

