## Supplementary text for "SoftWipe – a tool and benchmark to assess scientific software quality"

#### 1 Description of SoftWipe

`SoftWipe` is a pipeline, written in Python3, that uses predominantly freely available static and dynamic code analyzers to assess the code quality of software written in `C/C++`. `SoftWipe` initially compiles the software using the `clang` compiler (https://clang.llvm.org) and subsequently executes the software with *clang-sanitizers*. To accomplish this, the user is required to provide a text file with a command line call of the specific software. Subsequently, `SoftWipe` counts the lines of code, excluding comments and empty lines, and executes the static code analyzers described in Section 2. Using the output of the static code analyzers, `SoftWipe` computes a score for each static[1] analyzer and then outputs an overall score. `SoftWipe` retains all intermediate code analysis results such that the software developer gains an insight into the code quality and potential issues of his/her software. If one or several of the code analysis tools fail to produce a score, `SoftWipe` will exclude the respective category from the overall score calculation. The user is also able to exclude analysis tools from the overall score. This can be done via a respective command line switch (e.g., `--exclude-infer` to exclude the Infer analysis; see Section 2.7 for details).

[1]Strictly speaking, the clang-sanitizers are not static, but dynamic analyzers.

#### 2 Analysis tools

##### 2.1 Compiler and sanitizer

`SoftWipe` compiles the scientific software in our benchmark with the `clang` compiler with a large set of enabled warning flags. That is, it uses `-Weverything` with the exception of the warnings listed below which we chose to exclude as we considered them as being excessively pedantic.

- -Wno-padded
- -Wno-c++98-compat
- -Wno-c++98-pedantic
- -Wno-c++11-extensions
- -Wno-c99-compat
- -Wno-newline-eof
- -Wno-source-uses-openmp

We chose `clang`, as we empirically found that it typically produces more warnings than other compilers such as `gcc`. Further, we classify the warnings subjectively into three categories: *could-fix*, *should-fix* and *must-fix* weighted by 1, 2, and 3, respectively. Here a higher weight implies a higher severity (e.g., implicit type conversions that may result in a loss of precision). The weight assignments for warnings are shown in table 1. The classification was conducted via an open discussion in the slack channel of our lab that currently comprises 8 members that all have formal training in pure computer science.

As already mentioned, we also execute the compiled executable with `clang` sanitizers (`AdressSanitizer` and `UndefinedBehaviorSanitizer`). To execute the software, we use minimal toy examples as provided by the authors of the software, if available. In cases, where no examples are available, we set them up to the best of our abilities using freely available public datasets.

We employ the following sanitizer flags:

- -g
- -fno-omit-frame-pointer
- -fsanitize=address
- -fsanitize-recover=address
- -fsanitize=undefined

All sanitizer warnings receive a weight of 3 by default. We sum over the weighted warnings of the compiler and sanitizer and calculate the *compiler and sanitizer rate* as the weighted sum per total number of lines of code *(LOC)*.

| **warning** | **weight** |
|---|---|
| -Wc99-extensions | 1 |
| -Wcast-qual | 1 |
| -Wchar-subscripts | 1 |
| -Wcomment | 1 |
| -Wcovered-switch-default | 1 |
| -Wdate-time | 1 |
| -Wdeprecated | 1 |
| -Wdeprecated-dynamic-exception-spec | 1 |
| -Wdeprecated-register | 1 |
| -Wdocumentation | 1 |
| -Wdocumentation-deprecated-sync | 1 |
| -Wdocumentation-unknown-command | 1 |
| -Wexit-time-destructors | 1 |
| -Wextra-semi | 1 |
| -Wglobal-constructors | 1 |
| -Wgnu-zero-variadic-macro-arguments | 1 |
| -Wmissing-declarations | 1 |
| -Wmissing-noreturn | 1 |
| -Wold-style-cast | 1 |
| -Woverloaded-virtual | 1 |
| -Wparentheses-equality | 1 |
| -Wreorder | 1 |
| -Wshadow-field | 1 |
| -Wshadow-field-in-constructor | 1 |
| -Wshadow-field-in-constructor-modified | 1 |
| -Wunknown-pragmas | 1 |
| -Wunreachable-code | 1 |
| -Wunused-exception-parameter | 1 |
| -Wunused-function | 1 |
| -Wunused-macros | 1 |
| -Wunused-template | 1 |
| -Wvarargs | 1 |
| -Wweak-vtables | 1 |
| -Wwritable-strings | 1 |
| -Wc11-extensions | 2 |
| -Wcast-align | 2 |
| -Wcomma | 2 |
| -Wdeprecated-declarations | 2 |
| -Wdouble-promotion | 2 |
| -Wempty-body | 2 |
| -Wexpansion-to-defined | 2 |
| -Wfor-loop-analysis | 2 |
| -Wformat | 2 |
| -Wformat-extra-args | 2 |

| | |
|---|---|
| -Wformat-nonliteral | 2 |
| -Wgnu-binary-literal | 2 |
| -Wimplicit-int | 2 |
| -Winvalid-source-encoding | 2 |
| -Wlanguage-extension-token | 2 |
| -Wlogical-not-parentheses | 2 |
| -Wlogical-op-parentheses | 2 |
| -Wmacro-redefined | 2 |
| -Wmissing-prototypes | 2 |
| -Wmissing-variable-declarations | 2 |
| -Wnested-anon-types | 2 |
| -Wnonnull | 2 |
| -Wparentheses | 2 |
| -Wpedantic | 2 |
| -Wreserved-id-macro | 2 |
| -Wshadow | 2 |
| -Wstrict-prototypes | 2 |
| -Wswitch-bool | 2 |
| -Wswitch-enum | 2 |
| -Wundefined-func-template | 2 |
| -Wunreachable-code-break | 2 |
| -Wunreachable-code-loop-increment | 2 |
| -Wunreachable-code-return | 2 |
| -Wunused-parameter | 2 |
| -Wunused-private-field | 2 |
| -Wunused-value | 2 |
| -Wunused-variable | 2 |
| -Wvexing-parse | 2 |
| -Wvla | 2 |
| -Wvla-extension | 2 |
| -Wzero-as-null-pointer-constant | 2 |
| -Wabsolute-value | 3 |
| -Wbad-function-cast | 3 |
| -Wconditional-uninitialized | 3 |
| -Wconstant-conversion | 3 |
| -Wconversion | 3 |
| -Wdelete-non-virtual-dtor | 3 |
| -Wfloat-conversion | 3 |
| -Wfloat-equal | 3 |
| -Wformat-security | 3 |
| -Wheader-hygiene | 3 |
| -Wimplicit-fallthrough | 3 |
| -Winfinite-recursion | 3 |
| -Wliteral-conversion | 3 |
| -Wmultichar | 3 |
| -Wnon-virtual-dtor | 3 |

| | |
|---|---|
| -Wnull-arithmetic | 3 |
| -Wnull-conversion | 3 |
| -Woverlength-strings | 3 |
| -Wpointer-bool-conversion | 3 |
| -Wpointer-sign | 3 |
| -Wreturn-type | 3 |
| -Wself-assign | 3 |
| -Wself-assign-field | 3 |
| -Wself-assign-overloaded | 3 |
| -Wself-move | 3 |
| -Wshift-sign-overflow | 3 |
| -Wshorten-64-to-32 | 3 |
| -Wsign-compare | 3 |
| -Wsign-conversion | 3 |
| -Wsometimes-uninitialized | 3 |
| -Wstatic-self-init | 3 |
| -Wstring-plus-int | 3 |
| -Wstring-compare | 3 |
| -Wstring-conversion | 3 |
| -Wtautological-constant-compare | 3 |
| -Wtautological-pointer-compare | 3 |
| -Wtautological-type-limit-compare | 3 |
| -Wtautological-unsigned-zero-compare | 3 |
| -Wundef | 3 |
| -Wuninitialized | 3 |
| -Wvector-conversion | 3 |
| -Wabsolute-value | 3 |
| -Wbad-function-cast | 3 |

Table 1: Compiler warning weights

##### 2.2 Assertions

Assertions provide a means to ensure that conditions for a correct execution of a program are met. A study by Casalnuovo et al. [1] suggests that functions *with* assertions *do* have fewer defects. Thus, we consider a software containing more assertions as being of higher quality. `SoftWipe` outputs the *assertion rate* as the number of assertions (C-style `assert()`, `static_assert()`, or custom assert macros, if available) per total LOC. If the analyzed code contains custom assert macros, the user can specify them via regular expression (regex) using the following `SoftWipe` option `-a custom_assert_regex`.

##### 2.3 Cppcheck

`Cppcheck` (http://cppcheck.sourceforge.net) is a further static code analysis tool that detects undefined behavior and potentially dangerous coding constructs.

| warning | weight |
|---|---|
| information | 0 |
| style | 1 |
| performance | 1 |
| portability | 3 |
| error | 3 |
| warning | 3 |

Table 2: Cppcheck warning weights

We weight the warnings according to the weightings presented in Table 2 and output a *cppcheck rate* of the weighted warnings per total LOC.

#### 2.4 Lizard

`Lizard` (https://github.com/terryyin/lizard) is a cyclomatic complexity analyzer that outputs three metrics.

The first output is the 'cyclomatic complexity', which is a software metric to quantify the complexity/modularity of a program using the number of linearly independent paths in the control flow graph of the program. It assumes that intuitive complexity and graph-theoretic complexities correlate [2]. A linearly independent path is a path that has at least one unique edge that is not part of any other path. `Lizard` computes the cyclomatic complexity for each function, as well as the average cyclomatic complexity over all functions.

The second metric it calculates is the 'number of complex functions'. Lizard considers functions as being overly complex if their cyclomatic complexity, length, or parameter number exceeds a given threshold (i.e., 15 for cyclomatic complexity, 1000 for length, 100 for the number of parameters). The respective rate is calculated as the number of overly complex functions per total number of functions. As it is usually advisable to keep functions as short and simple as possible, these thresholds alarm the programmer once he exceeds them.

The third metric calculated by `Lizard` is the *unique rate* which outputs a score depending on the amount of duplicated code.

#### 2.5 KWStyle

For larger software projects it is common to enforce coding guidelines to produce more readable and maintainable code.

The `KWStyle` tool (https://kitware.github.io/KWStyle/) automatically checks for adherence to certain common coding style guidelines. It also allows the programmer to set up a set of custom guidelines in a `KWStyle.xml` file. `KWStyle` then analyses the code, detects violations of these guidelines and outputs respective warnings. At present, `SoftWipe` employs a general `KWStyle.xml` file which limits the line length, requires spaces between operators, and prohibits

| warning | weight |
|---|---|
| performance | 1 |
| readability | 1 |
| boost | 1 |
| cpp-core-guidelines | 1 |
| misc | 1 |
| modernize | 1 |
| bugprone | 2 |
| clang-analyzer | 2 |
| mpi | 2 |

Table 3: Clang-Tidy warning weights

| warning | weight |
|---|---|
| dead store | 1 |
| empty vector access | 3 |
| null dereference | 3 |
| memory leak | 3 |
| resource leak | 3 |
| uninitialized value | 3 |

Table 4: Infer warning weights

multiple statements in the same line. We calculate a `KWStyle` score depending on the warnings generated per total LOC.

#### 2.6 Clang-Tidy

`Clang-tidy` (https://clang.llvm.org/extra/clang-tidy/) is a static analysis tool which focuses on detecting typical programming errors, such as coding style violations or interface misuse which could lead to bugs. It generates several warning categories which we weight as shown in Table 3. We calculate the `Clang-tidy` rate as weighted warnings per LOC.

#### 2.7 Infer

`Infer` (https://github.com/facebook/infer) also is a static analysis tool that can detect severe issues such as memory leaks. We therefore weight most of the warnings produced by `infer` with a weight of 3, as shown in Table 4. As before, we calculate the `infer` rate as weighted warnings per LOC.

### 3 Scoring

Each of the analysis methods listed above returns a *rate* which we use to calculate the intermediate score (one score for each method) which is in turn used to calculate the *overall score* as an unweighted average over all intermediate scores. For the calculation of the intermediate scores we have two different approaches which we describe in the following. In order to make different analysis tools comparable, we calculate *worst* and *best* bounds from the corpus of tools that we have tested so far, excluding outliers. We call the rate $r$ of a tool an outlier, if it lies outside Tukey's fences[3], that is, for our set $R_c$, which contains all rates of programs in the benchmark for a category $c$, $r$ is an outlier *iff* $r \notin [Q_1 - k(Q_3 - Q_1), Q_3 + k(Q_3 - Q_1)]$, for upper and lower quartiles $Q_1, Q_3$ of $R_c$ and a non-negative constant $k$. We set $k := 1.5$, as proposed by Tukey. For analysis tools where a global best or worst bound exists (e.g., there can be no better case than observing 0 warnings), we explicitly set the bound to this value.

#### 3.1 Relative scoring

We use relative scoring to compare a set of software programs with each other. To achieve this, we employ a linear formula scaled between the worst and best bound such that a rate $x$ receives 0/10 if $x = \textit{worst}$ and 10/10 of $x = \textit{best}$. We update the worst and best bounds after each newly included software to the benchmark. Consequently, we have to update the relative scores in the benchmark every time we add new software. Hence, relative `SoftWipe` scores will not be stable over time as more software is being added and can not be easily referenced.

#### 3.2 Absolute fixed scoring

The absolute fixed score is the score which the user receives as feedback including the intermediate and overall results. This absolute fixed score does not change over time with the addition of new software to the benchmark and can hence be referenced in submission or publications. To achieve this, it does not suffice to simply fix the [*worst*, *best*] interval, since a rate $x$ (see Section 2 for detailed definitions of the rates) of the new software being added can exceed that interval resulting in a score below 0 or over 10. Fixing the maximum/minimum scores for rates that exceed the interval yields distinguishing among these rates impossible. Thus, we employ the Sigmoid function $\frac{1}{1+e^{-k(x-x_0)}}$ as the scoring function and scale the parameters $k$ and $x$ using the *curve_fit()* function from the *scipy* library to produce a score between 0 and 10. We use the following code to scale the Sigmoid:

```
d = best - worst
x = rate - worst
thresh = 0.90
```

```
xval = [(1 − thresh) * d, 0.25 * d, 0.5 * d, 0.75 * d, thresh * d]
yval = [(1 − thresh), 0.25, 0.5, 0.75, thresh]
popt, pcov = curve_fit(sigmoid, xval, yval)
return sigmoid(x, *popt)
```

For analysis methods where we can fix the worst or best boundaries a priori, we split the scoring function into a linear part and a Sigmoid part. In the case that we only fixed *worst*, the linear part is used for scores in $[0, 5]$ and sigmoid for scores are used in $(5, 10]$. The case where only *best* is fixed is implemented analogously.

We chose this specific approach due to its high rank correlation of 0.9972 to the relative score.

### 4 Executing SoftWipe

We developed and tested `SoftWipe` mainly on (and for) Linux-based systems. Thus the following recommendations may or may not work in the same way for Windows or MacOS. `SoftWipe` requires `Python3` and the following python packages:

- numpy $\geq$ 1.17.4
- scipy $\geq$ 1.3.3

as well as the following tools:

- Clang (https://clang.llvm.org)
- Cppcheck (http://cppcheck.sourceforge.net)
- Clang-Tidy (part of LLVM tools http://llvm.org)
- Lizard (https://github.com/terryyin/lizard)
- KWStyle (https://kitware.github.io/KWStyle/)
- Infer (https://github.com/facebook/infer)

For Debian-based systems `SoftWipe` can try to download the required tools automatically.
`SoftWipe` can be executed using the following command:

```
softwipe.py [-c | -C] [-m | -M | -l target [target ...]]
            [-e executefile] programdir
```

Where
**-c** is for software written in C, **-C** for software written in C++
**-m** is for Make-based builds, **-M** for CMake-based builds.
The makefile needs to use common variable names (e.g. ${CC} for the compiler, ${CFLAGS} for the compiler flags and ${LDFLAGS} for the linker flags) as

`SoftWipe` uses them to inject its own compilers and flags.
**-e** specifies a file that contains the shell command required to run the tested software, for instance,

```
./executable -arg1 -arg2
```

**programdir** specifies the root of the tested software
Further `SoftWipe` options are displayed via the `--help` command.

### 5 Benchmark

We used `SoftWipe` to analyze 51 software tools written in C/C++. We selected 20 of those tools by scrutinizing recent *Bioinformatics Application Notes* papers. We did not include GUI applications, pipelines, and software that we did not manage to execute. We did not analyze external libraries used by the softwares as they are frequently larger than the actual newly written software. This might lead to a distortion of the overall score. On the other hand one would typically not expect an application developer to fix the issues of the external libraries he/she is using. Libraries also tend to fill a certain niche which in general makes it unlikely to find suitable alternatives for every use case. When available, we used the execution examples provided by the tested softwares for sanitizer execution. For tools which did not include examples, we prepared simple execution examples ourselves. The results are available on the `SoftWipe github` page (https://github.com/adrianzap/softwipe/wiki/Code-Quality-Benchmark). We split the benchmark into two tables: One containing absolute values used to calculate the scores and one containing the absolute scores for each category as well as the absolute and relative overall scores. 'N/A' (not available) entries indicate that `SoftWipe` could not produce a score for the specific category (e.g., because the respective analysis tool terminated with an error). `SoftWipe` automatically excludes such results from the overall score.

### 6 Quality of Softwipe Code

We also assessed the quality of our `SoftWipe` code using the following commonly used static analyzers for python:

- Pylint (https://www.pylint.org/)
- Pyflakes (https://github.com/PyCQA/pyflakes)
- Radon (https://radon.readthedocs.io/en/latest/)

#### 6.1 Pylint

Pylint is a static analyzer that focuses on the adherence to Python's PEP 8 style guide (https://www.python.org/dev/peps/pep-0008/). It also checks for potential problems such as imports of deprecated modules. PEP8 is a widely known

Python style guide, recommended and written by the developer of Python Guido van Rossum himself. `SoftWipe` received a Pylint score of 7.14/10. However, this is mainly due to recommendations that, based on our personal assessment, would reduce the readability of our code. The complete output of Pylint is part of the Appendix 8.1.

#### 6.2 Pyflakes

Pyflakes is a static analyzer that focuses on finding source code errors. It does not assess coding style. This analyzer did not detect any issues with our `SoftWipe` code.

#### 6.3 Radon

Radon computes multiple code metrics that quantify code quality. These metrics are: Cyclomatic Complexity (CC), Halstead Metrics, and the Maintainability Index. We explain the CC in Section 2.4. Radon computes an average CC of 4.27, which yields the score "A" (low rist, simple blocks). The Halstead Metrics assume that programs are built out of operators and operands and uses these to estimate the difficulty to understand the program, the effort and time needed for the implementation, and the number of bugs in the implementation. We provide the calculated values in Table 5. The Maintainability Index estimates the difficulty to maintain (i.e., support, change, adapt) the program. For its calculation it considers CC, lines of code, the Halstead Volume (part of Halstead Metrics) as well as the percentage of comment lines. We receive an "A" (very high maintainability) for all SoftWipe files.

| **file name** | **difficulty** | **effort** | **time to program** | **bugs** |
|---|---|---|---|---|
| automatic_tool_installation.py | 2.95 | 743 | 41 | 0.084 |
| calculate_score_table.py | 8.04 | 8427 | 468 | 0.35 |
| classifications.py | 0 | 0 | 0 | 0 |
| compare_results.py | 9.9 | 9833 | 546 | 0.331 |
| compile_phase.py | 8.4 | 6810 | 378 | 0.270 |
| execution_phase.py | 3.55 | 664 | 37 | 0.062 |
| output_classes.py | 5.16 | 5721 | 317 | 0.37 |
| recalculate_scores_from_table.py | 9.7 | 8737 | 485 | 0.3 |
| scoring.py | 9.5 | 15190 | 843 | 0.533 |
| softwipe.py | 5.47 | 4259 | 236 | 0.26 |
| static_analysis_phase.py | 7.29 | 17193 | 955 | 0.79 |
| strings.py | 0.67 | 521 | 28 | 0.26 |
| tools_info.py | 0 | 0 | 0 | 0 |
| util.py | 2.12 | 251 | 14 | 0.04 |

Table 5: Radon Halstead Metrics for SoftWipe

##### 6.4 Lizard

We also used Lizard (for a description see Section 2.4) to assess the code quality of SoftWipe, since Lizard is not limited to C/C++. Based on Lizard we computed absolute scores (see Section 3.2) for the cyclomatic complexity, the Lizard warnings, and the unique code. The scores are 8.3/10, 8.1/10, and 9.9/10 respectively. The complete output is part of the Appendix 8.2.

#### 7 Future Improvements

While the `Infer` tool is fast to execute, we have noticed that it generally fails too often on some of the tools. To this end, we also plan to include `valgrind` in future releases of `SoftWipe` despite the fact that it is substantially slower. Another issue that we will address is to introduce version control for the code analysis tools that are being used by `SoftWipe`. For instance, versions 11 and 12 of `clang-tidy` tend to report more warnings than version 10.

### 8 Appendix

#### 8.1 Pylint output

The output of Pylint is available at
https://github.com/adrianzap/softwipe/blob/master/softwipe_code_quality/pylint_results.txt.

#### 8.2 Lizard output

The output of Lizard is available at
https://github.com/adrianzap/softwipe/blob/master/softwipe_code_quality/lizard_results.txt.
